# GO2Sum: Generating Human Readable Functional Summary of Proteins from GO Terms

**DOI:** 10.1101/2023.11.10.566665

**Authors:** Swagarika Jaharlal Giri, Nabil Ibtehaz, Daisuke Kihara

**Affiliations:** Department of Computer Science, Purdue University, West Lafayette, IN, United States; Department of Biological Sciences, Purdue University, West Lafayette, IN, United States

**Keywords:** protein function, protein function prediction, gene ontology, language model, summarizer

## Abstract

Understanding the biological functions of proteins is of fundamental importance in modern biology. To represent function of proteins, Gene Ontology (GO), a controlled vocabulary, is frequently used, because it is easy to handle by computer programs avoiding open-ended text interpretation. Particularly, the majority of current protein function prediction methods rely on GO terms. However, the extensive list of GO terms that describe a protein function can pose challenges for biologists when it comes to interpretation. In response to this issue, we developed GO2Sum (Gene Ontology terms Summarizer), a model that takes a set of GO terms as input and generates a human-readable summary using the T5 large language model. GO2Sum was developed by fine-tuning T5 on GO term assignments and free-text function descriptions for UniProt entries, enabling it to recreate function descriptions by concatenating GO term descriptions. Our results demonstrated that GO2Sum significantly outperforms the original T5 model that was trained on the entire web corpus in generating Function, Subunit Structure, and Pathway paragraphs for UniProt entries.

## Introduction

Elucidating the function of proteins is one of the most essential and important tasks in molecular biology, biochemistry, genetics, as well as bioinformatics. Conventionally, function of proteins is described in free text, such as those which we see in public protein and gene databases^1,2^ as it is natural and versatile for biologists to describe various aspects of functions and behaviors of proteins. However, a drawback of text representation is that it is challenging for computer programs to extract and use function information in various tasks such as finding proteins with a particular function from a genome or comparing functions of different proteins and to define functional similarity. To achieve machine-readable unified function description, Gene Ontology (GO) ^3,4^ a structured vocabulary for describing protein functions, has been developed about two decades ago, which is now well-established. GO classifies protein functions in three primary categories: Biological Process (BP), which pertains to pathway information; Cellular Component (CC), describing subcellular locations; and Molecular Function (MF), focusing on biochemical reactions of proteins. Using GO, the function of a protein can be represented as a list of GO terms. As each functional term is tokenized, GO has significantly facilitated computational studies of protein functions, such as quantitative comparison of protein functions and GO enrichment analysis^5^, which is indispensable for proteomics and genomics studies.

Protein function prediction is one of the research areas which has substantially benefit from GO^6^. Although the function of a protein needs to be ultimately determined by wet-lab experiments, computational prediction serves as a valuable tool by providing hypotheses and guiding biologists in designing experiments. Conventionally, computational function annotation (prediction) uses sequence database search ^7,8^ as a source of function information. Methods were developed that use sequence information with more thorough function information mining techniques ^9–12^. Other information used include protein domain composition in protein sequences ^13,14^, protein tertiary structures ^15^, protein networks ^16^, literature^16–19^, and combinations of multiple sources ^19–21^. The progress of computational function prediction has been objectively monitored by the community-wide function prediction assessment, the Critical Assessment of Function Annotation (CAFA) ^22^.

Most of the contemporary protein function prediction methods rely on GO. An output of protein function prediction comprises a list of GO terms, which can often be long, sometimes exceeding several dozen terms, and difficult for researchers to comprehend. In fact, the presentation of a list of GO terms has frequently led to questions and confusion among users of our protein function prediction web servers^23^.

In this study, our primary objective is to transform a list of GO terms into human-readable text, utilizing a recent advancement in Natural Language Processing (NLP) techniques. NLP, a field within computer science, has experienced significant progress, thanks to deep learning technologies^24^. With the use of transformer models, particularly Large Language Models (LLMs), we can now perform a multitude of NLP tasks, including translation, summarization, and question-answering, at a practical and efficient level.^25^. The work we present here, GO2Sum, takes as its input short text descriptions of a set of GO terms that annotate a protein and produces a text summary describing the protein’s function. To achieve this, we fine-tuned an LLM, the base version of Text-To-Text Transfer Transformer (T5) ^26^, on UniProt ^27^ entries with their GO annotations and functional descriptions.

LLMs have been increasingly used in bioinformatics domains. A pre-trained BERT language model ^28^ that was trained on the general text was further fine-tuned on biomedical literature to perform tasks such as sentence classification, dependency parsing ^29^, biomedical relation extraction, biomedical question answering ^30^, sentence similarity, and relation extraction^31^. PubMedBERT^32^ is a BERT model trained on PubMed abstracts and performs tasks including relation extraction, question answering, and document classification. Instead of BERT, a text-to-text transfer transformer model, T5^26^, as used in SciFive^33^, which was trained on biomedical corpora and performed language generation tasks such as relation extraction, natural language inference, and question-answering.

LLMs were also used for summarization tasks in the bioinformatics domain. Xie et al. used a knowledge infusion training framework to enhance multiple PLMs for summarizing biomedical literature^34^. Bidirectional and Auto-Regressive Transformers (BART)^35^ and the domain-aware pre-trained language model in BioBERTSum^36^ were used for abstractive and extractive summarization of biomedical evidence, respectively.

In GO2Sum, we employ T5 to summarize the functional descriptions from a list of GO terms. To the best of our knowledge, his work represents the pioneering effort in performing summarization within the protein function prediction domain. When selecting a technique for this general summarization task, we opted for T5 due to its superior performance demonstrated by the encoder-decoder-based architecture, outperforming earlier techniques.^26,37,38^. From UniProt entries, we extracted three functional descriptions, Function, Subunit Structure, and Pathway. For each of these descriptions, we trained a separate T5 model. We showed that the fine-tunned T5 model performed significantly better than the pre-trained vanilla T5 in reproducing these function descriptions when we evaluated with three embedding-based metrics, BERT^28^, MiniLM^39^, and BioSentVec^40^ as well as three sentence distance-based metrics, WMS^41^, SMS^41^, and W+SMS^41^. Moreover, in GO2Sum we provide a confidence level of output summary by considering the probability of multiple variations of summary texts in beam search. Finally, we applied GO2Sum to the predicted GO terms generated by Phylo-PFP^9^ and demonstrated that, in most cases, the summaries produced by GO2Sum exhibit sufficient accuracy even when derived from predicted GO terms.

### Framework of GO2Sum

The GO2Sum workflow, illustrated in Fig. 1, begins with a set of input GO terms, each accompanied by a text description. For instance, GO:0000049 is associated with the description ‘Binding to a transfer RNA.’ These GO term descriptions are concatenated into a document, serving as the input for the summarizer model, T5. The summarizer then generates a paragraph that elucidates the function of the input protein.

**Figure 1.**
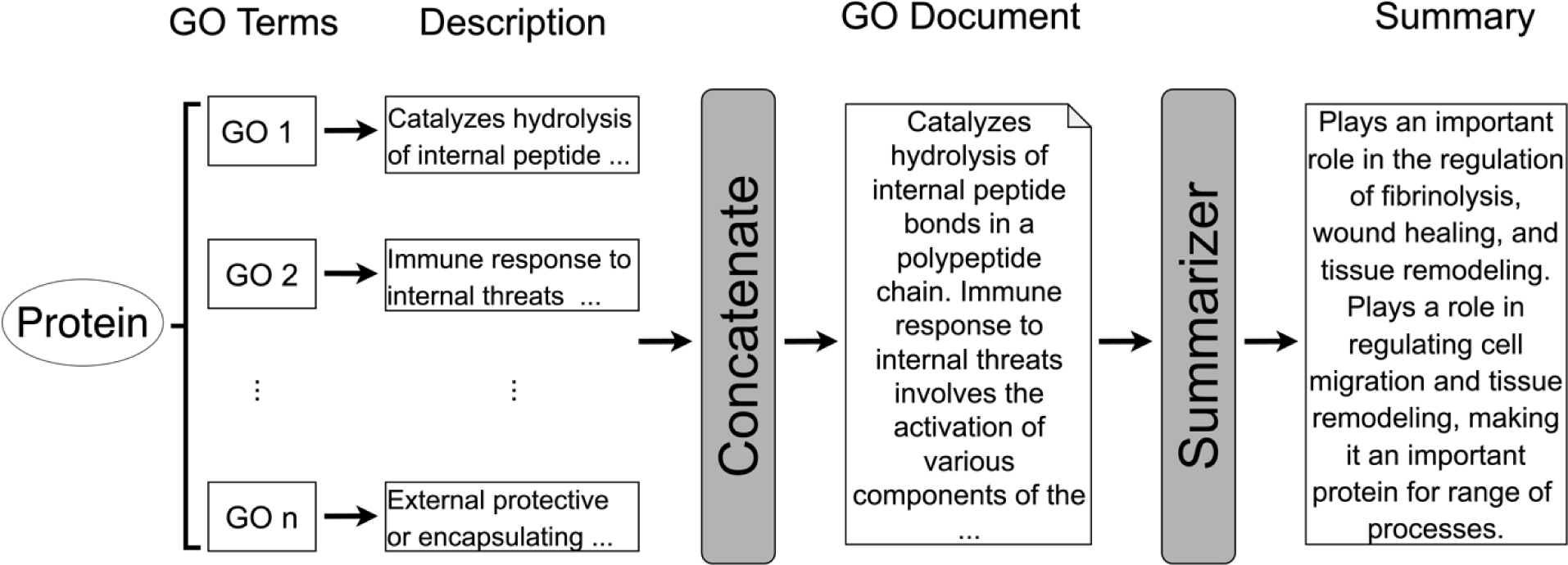
GO2Sum Workflow.

The dataset of GO terms and protein text descriptions was sourced from SwissProt (Release: February 11, 2022). Out of the 542,953 proteins included in this SwissProt release, we selected 518,422 proteins that had at least one GO term annotation. For each GO term, we obtained the text description from the Gene Ontology Consortium (Release: November 16, 2021). The length of these GO term descriptions ranged from 3 to 7,836 words, with an average of 310 words. Meanwhile, we gathered three different paragraphs that described the function of each protein from their SwissProt entries. These paragraphs included the general function description, usually presented at the top of the UniProt entry (referred to as “Function in UniProt”). We also collected two additional paragraphs: one related to “Subunit Structure”, which details molecular interactions with other proteins, and the other providing “Pathway Information”, explaining the metabolic pathways associated with the protein. These three paragraphs served as the ground truth for the protein’s function description. For reference, some examples of these three function description paragraphs can be found in Supplementary Information 1.

To reduce redundancy of proteins of similar function, we filtered out proteins that had 90% or more identical GO term annotations. This process reduced the dataset to 109,658 proteins from 542,953. As an entry often does not have all three functional descriptions of Function, Subunit, and Pathway, we constructed three separate datasets for each. The dataset for Function, SubUnit and Pathway had 97,600, 62,340, 14,600 proteins, respectively.

Following earlier studies of text summarization^42,43^ we opted for an 80/10/10 split for training/validation/testing. Therefore, the training/validation/test sets of the Function dataset included 78,080/9,760/9,760 entries, Pathway had 11,680/1,460/1,460 entries, and Subunit had 49,872/6,234/6,234 entries, respectively. Due to the limitation of the size of GPU memory we used, up to 1,024 tokens or words were used from a concatenated GO document if it exceeded the size.

Among several T5 models with different parameter sizes, we used the T5 base model with 220 million parameters, 12 transformer layers, and 12 attention heads. Computation of training was performed on NVIDIA RTX5500 24GB memory and inference was performed on NVIDIA RTX A6000 GPU with 48GB memory. We trained three models, each for producing paragraphs for Function, Subunit structure, and Pathway sections in UniProt, respectively. Supplementary Fig. 1 shows how loss changed during the training process of the models. For the loss function, token-level cross-entropy loss between the predicted summary and the ground truth summary was used.

### GO Summary for UniProt Function

First, we assess the summarization performance of GO2Sum for Function paragraphs in UniProt. We employ two types of scoring methods: embedding-based scores and sentence mover distance-based similarities. Embedding-based scores provide a quantitative comparison between the output generated by GO2Sum and the ground truth Function paragraphs obtained from UniProt entries. These scores are more suitable over traditional that rely on n-gram overlaps, such as ROUGE ^44^ and BLEU ^45^, as embedding take into account word meaning and semantic context. We used three different embedding models, BERT^28^, MiniLM^39^, and BioSentVec^40^ embeddings. BERT and MiniLM are pre-trained language models for general natural language processing (NLP) tasks while BioSentVec are specially designed for biomedical texts. Using these embeddings, paragraph similarity is quantified through cosine similarity with a score range from -1.0 (complete irrelevance) to 1.0 (perfect match).

For the second type of scores, mover distance-based similarity, we used three metrics, Word Mover’s Similarity (WMS) ^41^, Sentence Mover’s Similarity (SMS) ^41^, and Sentence and Word Mover’s Similarity (S+WMS) ^41^. WMS combines bag-of-word histogram representations with word embedding similarity. SMS treats documents as bags of sentence embeddings and solves a linear optimization problem. S+WMS represents each document as a collection of words and sentences, solving the same linear equation as WMS. These similarity scores, produced by SMS, WMS, and S+WMS, are relative measures of similarity between two documents. They fall within the range of 0 to 1 and indicate the minimum cost required to transform one document into the other.

In Fig. 2a, 2b, and 2c, we compared the performance of GO2Sum relative to the vanilla T5 in reproducing UniProt Function. Across all three embedding-based scores, GO2Sum’s generated summaries achieved higher scores than vanilla T5 for the majority of UniProt entries. Specifically, GO2Sum’s summaries outperformed vanilla T5 for 95.5%, 96.7%, and 98.0% of UniProt entries with BERT, MiniLM, and BioSentVec embeddings, respectively. These scores are highly correlated, as illustrated in Fig. 2d, where the correlation between MiniLM and BioSentVec had a Pearson’s correlation coefficient of 0.886 (for additional score pairs, refer to Supplementary Fig. 2). Given this high correlation, we use the average score of the three for evaluation in the rest of the discussion.

**Figure 2.**
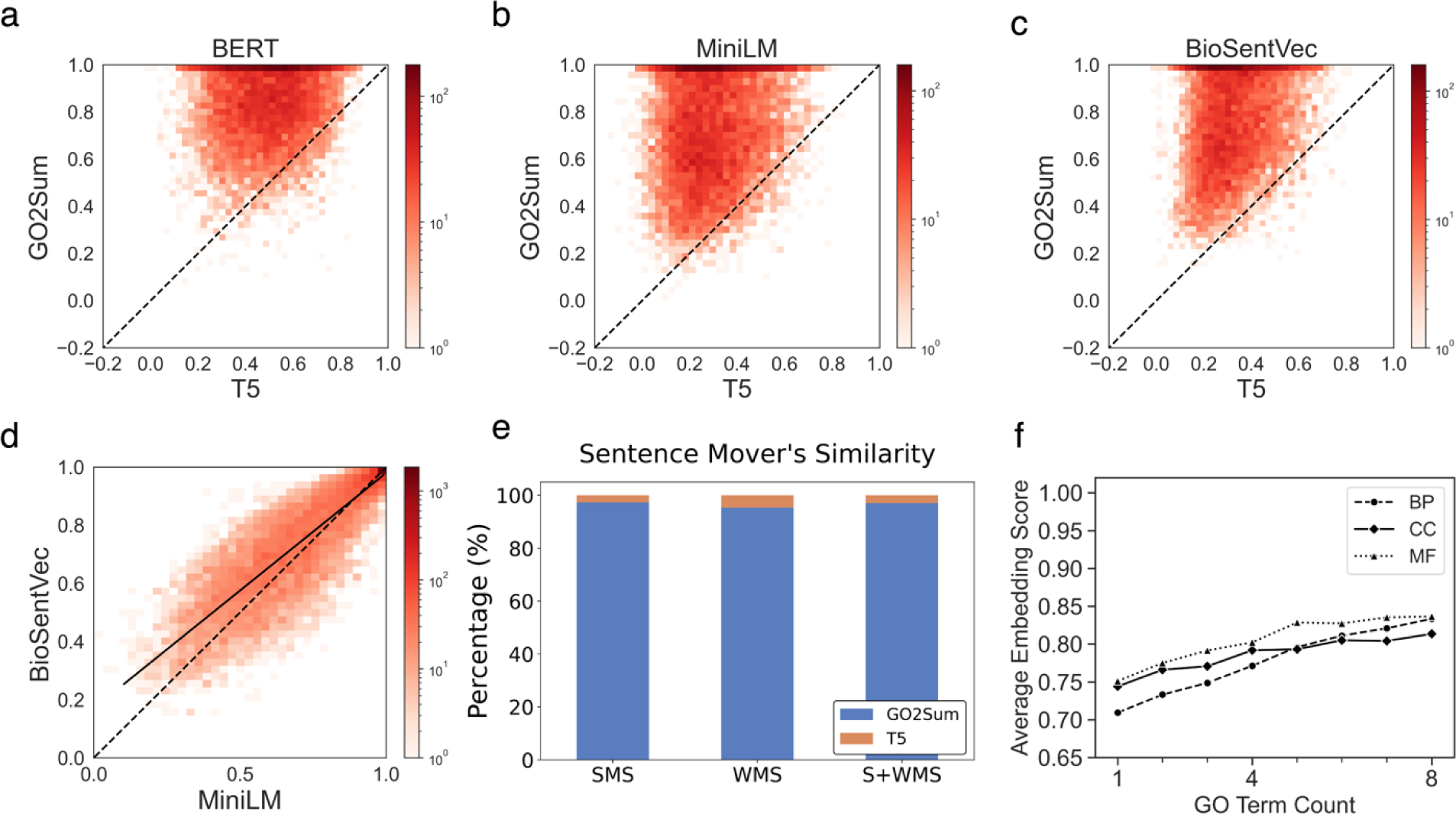
Evaluation scores for UniProt Function paragraphs. **a-c**, Comparison between GO2Sum and the vanilla T5 using the three embedding-basedℒscores, **a**, BERT, **b**, MiniLM, and **c**, BioSentVec. The number of UniProt entries used was 9,760. **d**, Comparison between MiniLM and BioSentVec. The correlation coefficient was 0.886. The dashed line is y=x line and the solid line is regression line. **e**, Comparison between GO2Sum and the vanilla T5 using the three sentence-mover similarity-based metrics, WMS, SMS, and S+WMS. The number of wins (i.e., entries where GO2Sum produced closer paragraph to ground truth than the vanilla T5) are shown in blue. **f**, impact of reducing GO terms. From a set of proteins, we reduced GO terms from their annotations randomly and observed how the average embedding score changed. The number of proteins in the dataset for BP, CC, and was 215, 130, and 55, respectively. The x-axis shows the number of remaining GO terms in GO annotations after removal of a certain number of GO terms. For each target, 1, 2, 3, …, and 7 GO terms were removed randomly three times and the embedding score was averaged over the three trials. Then, the value from each target was averaged across all the targets and plotted along the y-axis.

In Fig. 2e, we used the three mover-distance-based similarity scores to compare GO2Sum and vanilla T5. The results show that the summary by GO2Sum was more similar to the ground truth than vanilla T5 for almost all UniProt entries, 95.3%, 97.3%, and 97.0% of the cases by WMS, SMS, and S+WMS, respectively. When we took a close look at the remaining cases where GO2Sum is less similar to UniProt Function, typically those are cases that both GO2Sum and T5 failed to produce meaningful paragraphs. For example, The UniProt entry Q4R7M2 describes the protein as a dipeptidase that is capable of hydrolyzing cystinyl-bis-glycine, but unable to hydrolyze leukotriene D4 into leukotriene E4. Neither T5 nor GO2Sum was successful for this target. T5 produced paragraph “a phospholipid bilayer and associated proteins. a phospholipid bi,” while GO2Sum produced “Has a role in meiosis”, both of which were irrelevant to the actual function. The average embedding-based score of GO2Sum and T5 were 0.21 and 0.28 for such cases where GO2Sum lost over T5, indicating that summary of both GO2Sum and T5 were in a very poor quality.

To investigate the impact of the number of GO terms on GO2Sum’s Function summary generation, we conducted an experiment depicted in Fig. 2f. We systematically removed a fixed number of GO terms from the target protein’s GO annotation and examined the average of the three embedding scores. For this experiment, we curated a dataset of protein targets for each of the three GO categories. In the case of the Biological Process (BP) category, we selected proteins with exactly eight GO terms within their annotation, at least one GO term for each of the other categories (Cellular Component, CC, and Molecular Function, MF), and a total count of GO terms ranging from 10 to 24. This procedure yielded datasets of 215, 130, and 59 proteins for BP, CC, and MF, respectively. From these datasets, we randomly removed a fixed number of GO terms (ranging from 1 to 7) from each protein’s GO annotation. Subsequently, we ran GO2Sum with the remaining GO terms to generate Function summaries and evaluated the summaries using the average of the three embedding scores. To account for randomness, we repeated this experiment three times and averaged the results for each protein.

As shown in the graph, starting from the score of around 0.80 when there were eight GO terms (full annotation), the embedding score gradually declined as more GO terms were removed, ultimately reaching around 0.70 to 0.75 when only one GO term of that category remained. It’s worth noting that the score did not drop further because GO terms of other categories were retained throughout the removal process. This effect varied among different ontologies. Removing Biological Process (BP) terms had a more pronounced impact, resulting in a rapid decline in the embedding score. With an initial eight BP terms, the mean average embedding score was 0.83, decreasing to 0.70 when only one BP term remained. In contrast, Cellular Component (CC) terms had the least impact on Function annotation, with an average embedding score of 0.81 with all eight GO terms, decreasing to 0.74 when one GO term remained.

In Table 1, we present several examples of generated summaries. Each example includes a Function paragraph from UniProt, which is compared with the summaries produced by GO2Sum and the vanilla T5. The first three examples (UniProt ID: P0DMN7, Q86A79, and Q15438) showcase cases where GO2Sum achieved impressive average embedding-based scores ranging from 0.93 to 0.98. As demonstrated, with these high scores, GO2Sum generated summaries that closely matched the ground truth, with only negligible differences. These differences were typically minor, such as the absence of a specific protein name or slight variations in wording with the same meaning. In contrast, the vanilla T5 produced shorter, incomplete phrases, making the disparity between GO2Sum and T5 evident.

**Table1.**
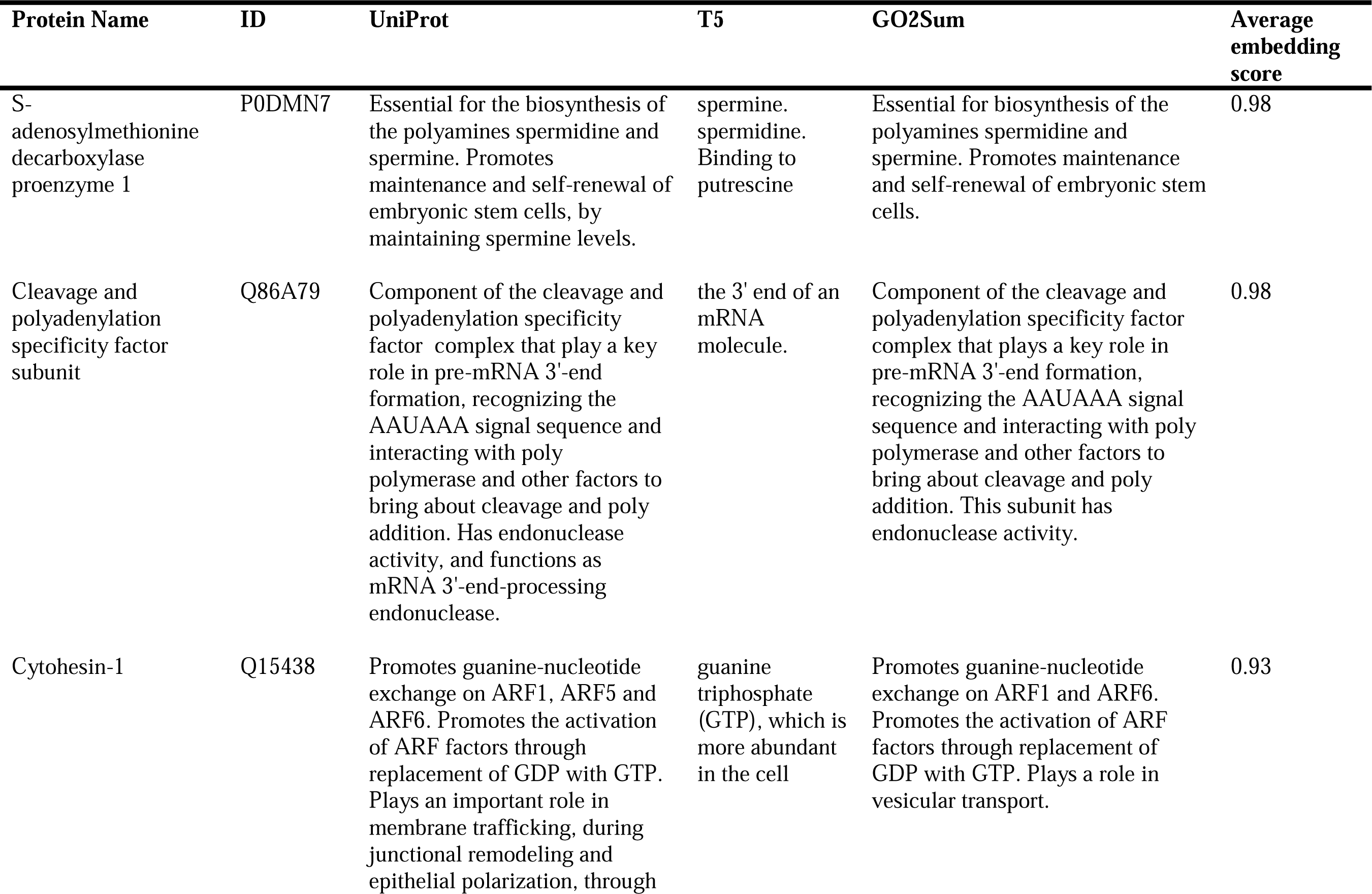
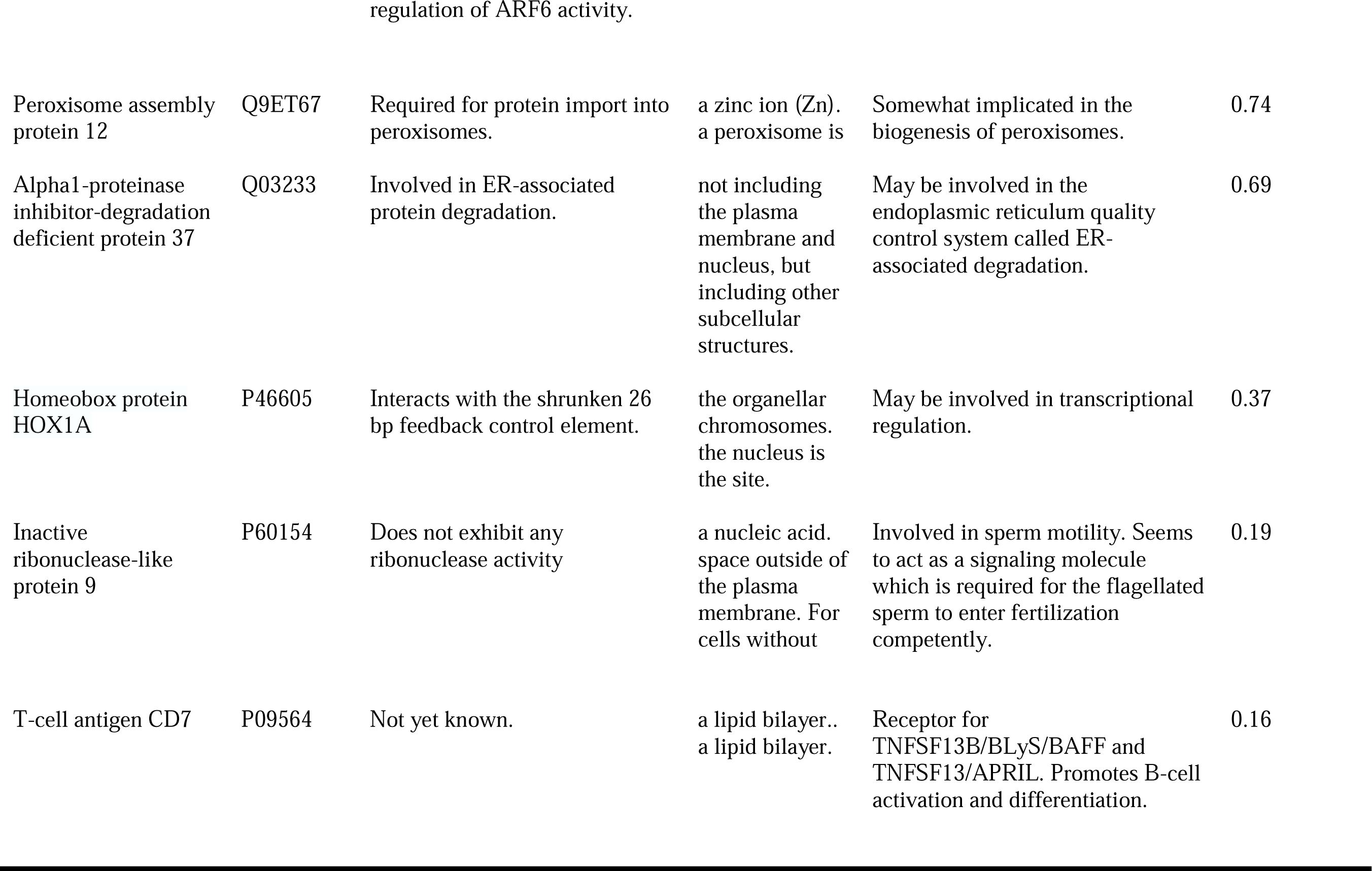
Examples of Summary generated for UniProt Function.

The next two examples, Q03233 and Q9ET67, are cases where GO2Sum had a moderate average embedding-based score of 0.69 to 0.74. These proteins have relatively brief function descriptions in UniProt. While GO2Sum’s summaries correctly capture the protein functions, there are differences in wording and expression compared to the UniProt descriptions. In contrast, the vanilla T5 model provides fragmented phrases. Notably, T5’s summary for Q03233 is incorrect as it fails to mention protein degradation.

For P46605 and P60154, GO2Sum’s summaries obtained low scores of 0.37 and 0.19, respectively. In the case of P46605, GO2Sum’s summary captures essentially correct information, although the ground truth provides more specific details, indicating its interaction with a particular DNA element. The next entry, Inactive ribonuclease-like protein 9 (P60154) is an interesting example, where GO2Sum has a low average embedding score of 0.19 indicating that the generated summary is dissimilar to the ground truth. For this entry, UniProt only added a short Function description that negates the ribonuclease activity despite the sequence similarity, which may not be comprehensive summary of the known functional activity of this protein. On the other hand, the GO document of this entry, i.e., the concatenated explanations of GO terms read “Binding to a nucleic acid. The space external to the outermost structure of a cell. For cells without external protective or external encapsulating structures, this refers to space outside of the plasma membrane. This term covers the host cell environment outside an intracellular parasite. The process in which the controlled movement of a flagellated sperm cell is initiated as part of the process required for flagellated sperm to reach fertilization competence.”. Thus, there are publications^46^ that indicates the involvement of sperm, specifically sperm mobility, from which these GO terms are assigned. Therefore, the summary generated by GO2Sum would actually be a better summary for the mentioned document as it accurately captures the main concepts related to the function of this protein, which is involved in sperm motility and acts as a signaling molecule necessary for the proper movement of spermatozoa.

The last example, T-cell antigen CD7 (P09564) has a similar story as the previous example. UniProt describes its function as “Not yet known”, which is obviously uninformative, despite that this protein has seven GO term annotations (GO:0016020, GO:0005886, GO:0038023, GO:0002250, GO:0006955, GO:0042110, GO:0007169). The summary by GO2Sum has a low average embedding-based score of 0.16. However, considering the annotated GO terms and references^47^ of this entry, we see that GO2Sum’s summary, “Receptor for TNFSF13B/BLyS/BAFF and TNFSF13/APRIL. Promotes B-cell activation and differentiation.” is accurate and more informative than UniProt Function description

### GO Summary for UniProt Subunit Structure

Next, we discuss summary for Subunit structure section in UniProt. As shown in Supplementary Information 1, paragraphs of Subunit structure tend to be shorter compared to the Function paragraphs. The paragraphs contain information related to subunit structure, such as protein interactions (e.g., interaction with some protein), stoichiometry e.g., homotetramers and homotrimers. They often use specific protein names and IDs that can make it challenging for the models to produce it accurately.

Figure 3a illustrates the performance of GO2Sum in comparison to the vanilla T5, using the average of the three embedding-based scores. Similar to the findings in the Function section reported in Figure 2, GO2Sum achieved a higher score than T5 in 99.2% of the entries. The results for the three individual scores (BERT, MiniLM, and BioSentVec) are detailed in Supplementary Figure 3, and these scores exhibit a high correlation, as shown in Supplementary Fig. 3. In Figure 3b, we present an evaluation using mover distance-based similarity scores. GO2Sum’s summaries were found to be closer to the ground truth in UniProt entries compared to the vanilla T5 for 96.4% using Word Mover’s Similarity (WMS), 95.3% using Sentence Mover’s Similarity (SMS), and 96.8% using Sentence and Word Mover’s Similarity (S+WMS).

**Figure 3.**
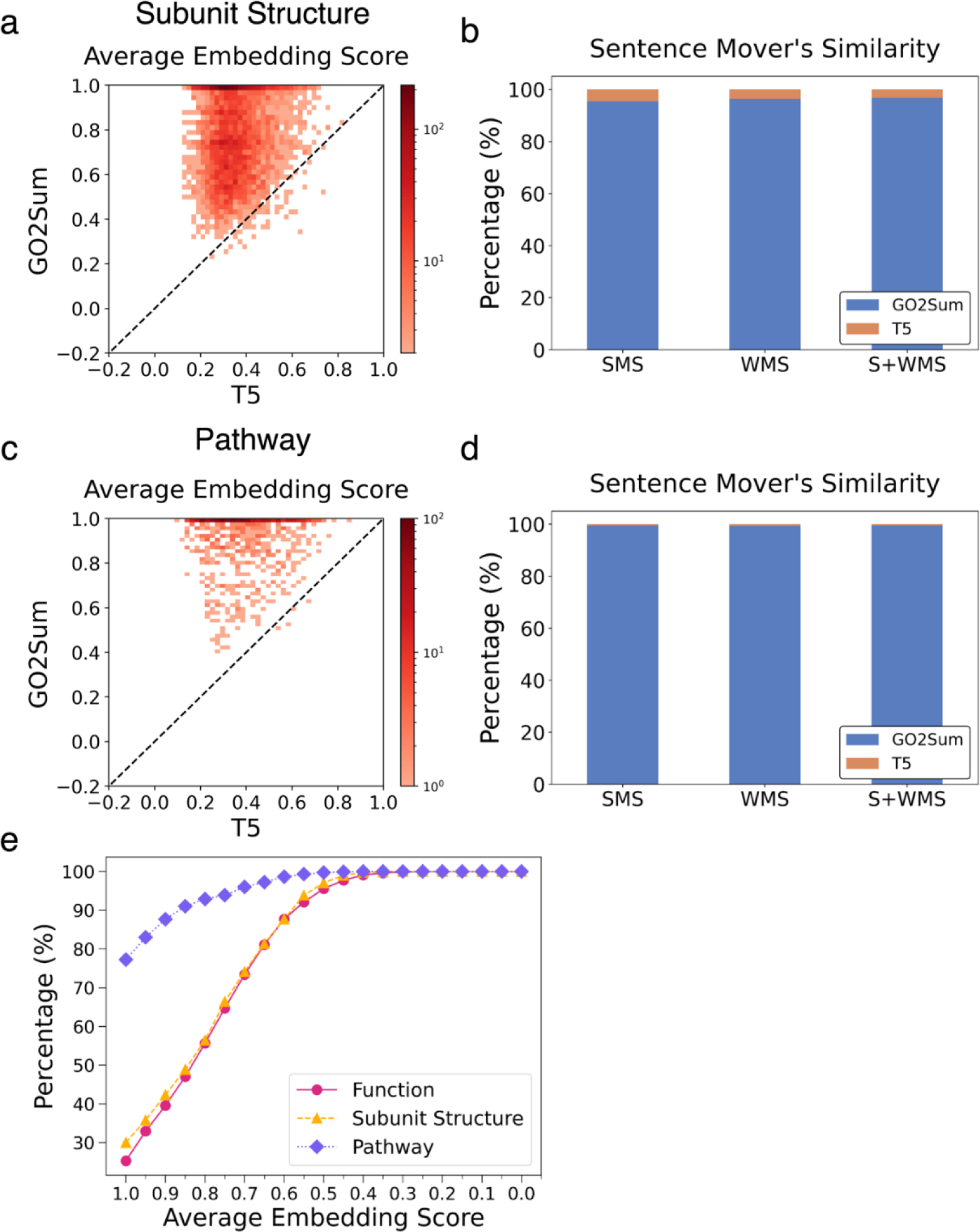
Comparison of GO2Sum and the vanilla T5 on the UniProt Subunit structure paragraphs and Pathway paragraphs. **a**, the average embedding score for Subunit structure paragraphs. GO2Sum outperforms vanilla T5 for 99.2% of cases. **b**, the Sentence mover’s similarity for Subunit structure paragraphs. The number of wins for GO2Sum for SMS, WMS and S+WMS is 95.3%, 96.4%, and 96.8% respectively. **c**, the average embedding score for Pathway paragraphs. GO2Sum outperform vanilla T5 for 99.8% of cases. **d**, the Sentence mover’s similarity for Pathway paragraphs. The number of wins for GO2Sum for SMS, WMS, and S+WMS was 99.5%, 99.3%, and 99.5%, respectively **e**, The Cumulative distribution of the average embedding score for Function, Subunit, and Pathway paragraphs.

In Supplementary Table 1, we provided examples of summaries generated by GO2Sum and T5. In some cases, GO2Sum’s summaries are generally in good agreement with the ground truth but lack specific protein names. This is attributed to the absence of such specific information in the GO descriptions.

### GO Summary for UniProt Pathway

The last UniProt sections to examine is Pathway descriptions. As shown in Fig. 3c and 3d, paragraphs made by GO2Sum clearly have a higher average embedding-score than the vanilla T5 (Fig. 3c) and also in terms of the Sentence Mover’s Similarity scores (Fig. 3d) for almost all the entries. Therefore, to summarize, consistently for all the three UniProt sections, Function, Subunit, and Pathway, GO2Sum made more meaningful and correct paragraphs. Examples of GO2Sum’s outputs are provided in Supplementary Table 2. Similar to examples shown in Supplementary Table 1 for Subunit structure, GO2Sum performed fundamentally better than the vanilla T5. Often the average embedding score of GO2Sum’s summary was low because it lacked specific information.

### Average embedding score distribution for the three UniProt sections

In Fig. 3e, we present the distribution of average embedding scores for GO2Sum’s outputs in the three UniProt sections to summarize its performance. Notably, the Pathway section exhibited a distinct score distribution, with 77.3% of entries achieving high scores between 1.0 and 0.95. In contrast, both Function and Subunit Structure summaries had only 25.3% and 30.0%, respectively, falling within the high score range. This divergence can be primarily attributed to paragraph length, with Pathway paragraphs being substantially shorter than those in the other two sections. On average, the word lengths for Pathway, Subunit, and Function summaries were 7.8, 28.4, and 56.1, respectively. Longer paragraphs, in general, tend to be more complex and challenging to reproduce accurately than shorter ones.

### Human Evaluation

To validate the embedding scores used in our work, we conducted a comparison with evaluations by human biology experts. We focused on assessing how the evaluations by human experts correlate with the embedding scores, particularly when the embedding score indicates high quality, i.e., favorable GO2Sum outputs. We engaged six human biology experts, four of whom hold master’s degrees in biology, while the other two possess bachelor’s degrees in biology or related fields such as biotechnology. The human evaluators assigned scores between 1 and 4, with 4 indicating the highest quality for each GO2Sum output.

We prepared two sets of protein targets for evaluation. The first set included 100 proteins, evenly distributed based on their average embedding scores. We randomly selected 10 proteins for each 0.1 interval, ranging from 0.1 to 0.2, 0.2 to 0.3, and so on. The second set comprised 20 randomly selected proteins with high average embedding scores between 0.8 and 1.0. We provided human experts with instructions, as detailed in Supplementary Information 2. In Fig. 4a, we present the Pearson correlation coefficients between human evaluators. Evaluator 3 exhibited a moderate correlation, ranging from 0.32 to 0.45, with other evaluators. On the other hand, the remaining five evaluators demonstrated strong agreement, with correlations ranging from 0.57 to 0.85.

**Figure 4.**
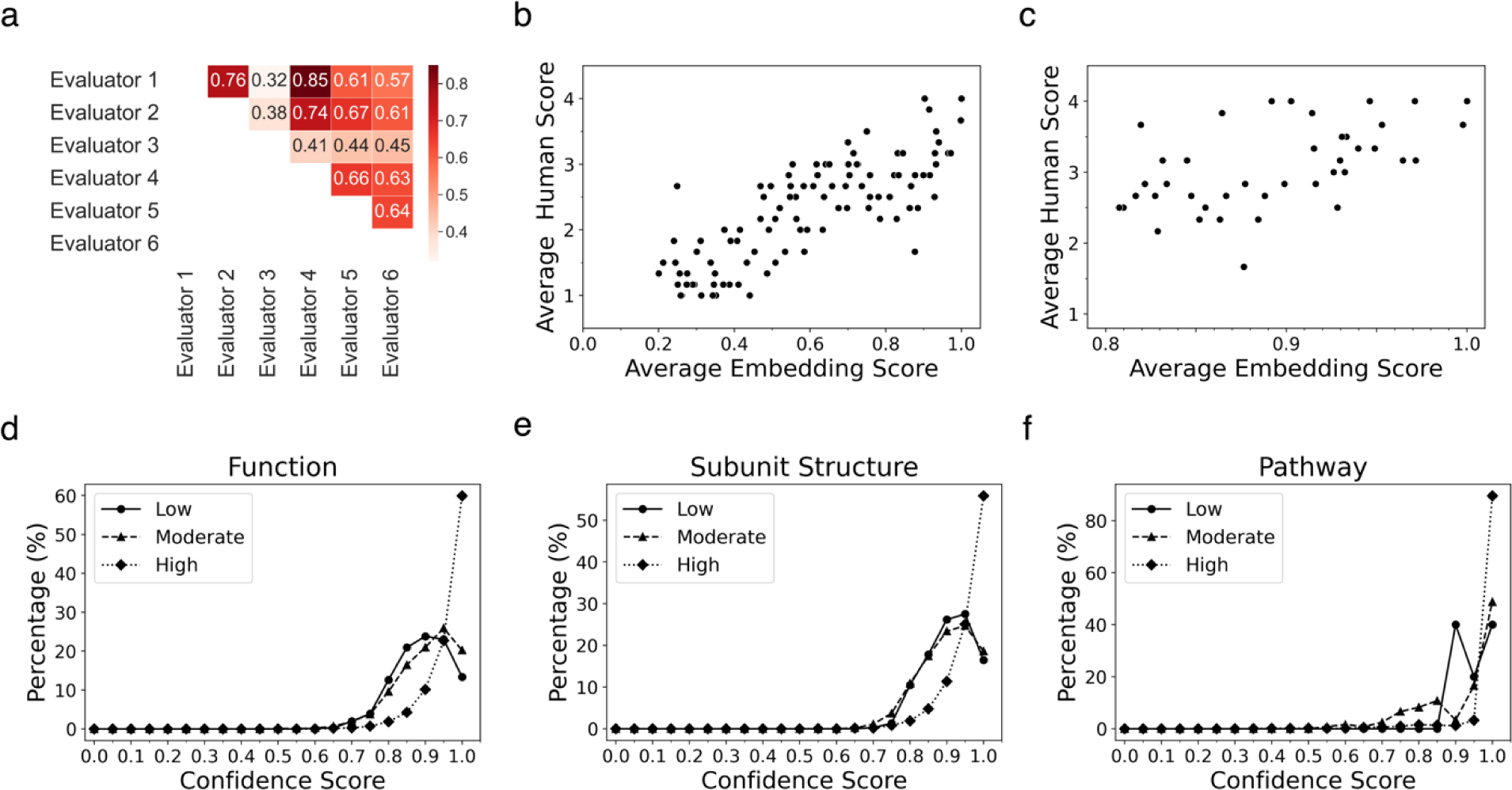
Expert human evaluation and the confidence score of paragraphs generated by GO2Sum. **a**, Pearson Correlation Coefficient between pairs of human evaluators for the dataset of 100 evenly distributed entries. **b**, Comparison between the average embedding score and the average of human evaluators score. The score correlation for the 100 proteins entry dataset that have the average embedding score between 0.1 to 1.0. **c**, the second dataset of 45 protein entries with a high average embedding score between 0.8 to 1.0. **d**, Distribution of the confidence score for entries classified into three classes based on the average embedding score. High, [0.8, 1.0]; moderate, [0.5, 0.8); low, [0, 0.5). Function paragraphs. 4,591, 4,400, and 769 proteins were included in the high, moderate, and low score class, respectively. **e**, Subunit Structure paragraphs. 3,046, 2,806, and 382 in high, moderate, and low. **f**, Pathway paragraphs. 1,329, 121, and 10 proteins in high, moderate, and low.

In Fig. 4b-c, we assessed the agreement between the average embedding score we used and the evaluations by human experts. The y-axis in Fig. 4b-c represent the average score assigned by the six human evaluators. As shown in Fig. 4a, the average embedding score demonstrates a strong correlation with the assessments made by human experts, with a Pearson correlation coefficient of 0.804. In Fig. 4b, we focused on 45 proteins with high average embedding scores ranging from 0.8 to 1.0. This selection included 25 high-scoring entries from dataset 1 and 20 proteins from dataset 2. Nearly all entries received an average human evaluator score of 2 or higher, with an overall average of 3.15. When considering entries with an even higher embedding score range of 0.9 to 1.0, the average human score rose to 3.52. Consequently, we can conclude that, on the whole, the average embedding score aligns sufficiently with evaluations by human biologists. Notably, high-scoring summaries generated by GO2Sum are highly reliable, as confirmed by human evaluators.

Although the embedding score agrees well with human evaluation, we observed two outliers where human evaluation differed from the embedding score. In the first case, Protein A3DNG9, a ribosomal RNA small subunit methyltransferase (Nep1), had a UniProt Function description: ‘Methyltransferase involved in ribosomal biogenesis. Specifically catalyzes the N1-methylation of the pseudouridine corresponding to position 914 in *M. jannaschii* 16S rRNA.’ GO2Sum summarized the GO terms as: ‘Methyltransferase involved in ribosomal biogenesis. Specifically catalyzes the N1-methylation of pseudouridine at position 967 in 16S rRNA. Is not able to methylate uridine at this position.’ Despite the high average embedding score of 0.87, human evaluators assigned an average score of 1.66 because the last sentence contradicted the preceding sentences.

Another example, O74523 (Putative ATPase inhibitor, mitochondrial), presents an opposite case, where the average embedding score was low (0.49), yet human evaluation scored it high at 2.66. The GO2Sum’s output, ‘Thought to be a regulatory component of the mitochondrial ATP-synthesizing complex in the mitochondria,’ lacked a term indicating that the protein ‘inhibits.’ However, aside from this omission, the overall function was accurate.

### Confidence score for summary outputs by GO2Sum

As T5 generates a paragraph using beam search, each generated paragraph is associated with a probability value. These probability values enable us to compute a score that reflects the confidence or dominance of a paragraph compared to alternative paragraphs explored during the generation process. The confidence score of a top-scoring paragraph *i* is defined as follows:

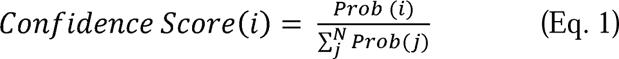

where *N* is the number of paragraphs generated at the last step of a beam search, which was set to 4 in this work.

In Fig. 4d-f, we present the distribution of confidence scores for UniProt summaries, categorized into three classes based on accuracy determined by average embedding scores. The high-scoring class included proteins with an average embedding score between 0.8 to 1.0, the moderate class ranged from 0.5 up to 0.8, and the low class included scores below 0.5. The plots reveal a clear trend: highly accurate summaries are typically associated with high confidence scores. This trend holds true for all three types of summary paragraphs (Function, Subunit, and Pathway). For instance, in the Function paragraphs, 82.6% of proteins in the high-scoring class have a confidence score exceeding 0.9. The proportion of highly confident summaries with a score over 0.9 decreases to 45.0% for the moderate class and 36.4% for the low-scoring class.

### Applying GO2Sum for GO term predictions by Phylo-PFP

Finally, we applied GO2Sum to predicted GO terms obtained from Phyo-PFP^9^, one of highly accurate protein function prediction methods. Predicted GO terms may not perfectly represent the actual functions of proteins, posing a challenge to GO2Sum’s summarization. To assess GO2Sum’s utility in describing predicted protein functions, we conducted tests across a wide range of accuracy levels using a realistic scenario of GO predictions. For the test set of proteins, we selected common proteins shared between the test set used in this study and the test set from the work of ContactPFP^15^, encompassing a total of 9,642 proteins. Among these, 843, 116, and 632 proteins were shared for Function, Pathway, and Subunit Structure paragraphs, respectively.

In Fig. 5a, b, c, the average embedding score of summaries generated from predicted GO terms were plotted relative to results from the ground truth GO terms taken from UniProt. Naturally, summaries generated from predicted GO terms have a lower score for most of the cases (82.7% for Function; 77.1% for Subunit, and 94.8% for Pathway). However, it is noteworthy that the majority of summaries generated from predicted GO terms still maintain a moderate average embedding score of 0.5 or higher, the level which is practically accurate and useful for users. Specifically, for Function paragraphs, 73.7% achieved a score of 0.5 or higher, while for Subunit paragraphs, the figure was 79.9%, and for Pathway paragraphs, it was 95.7%.

**Figure 5.**
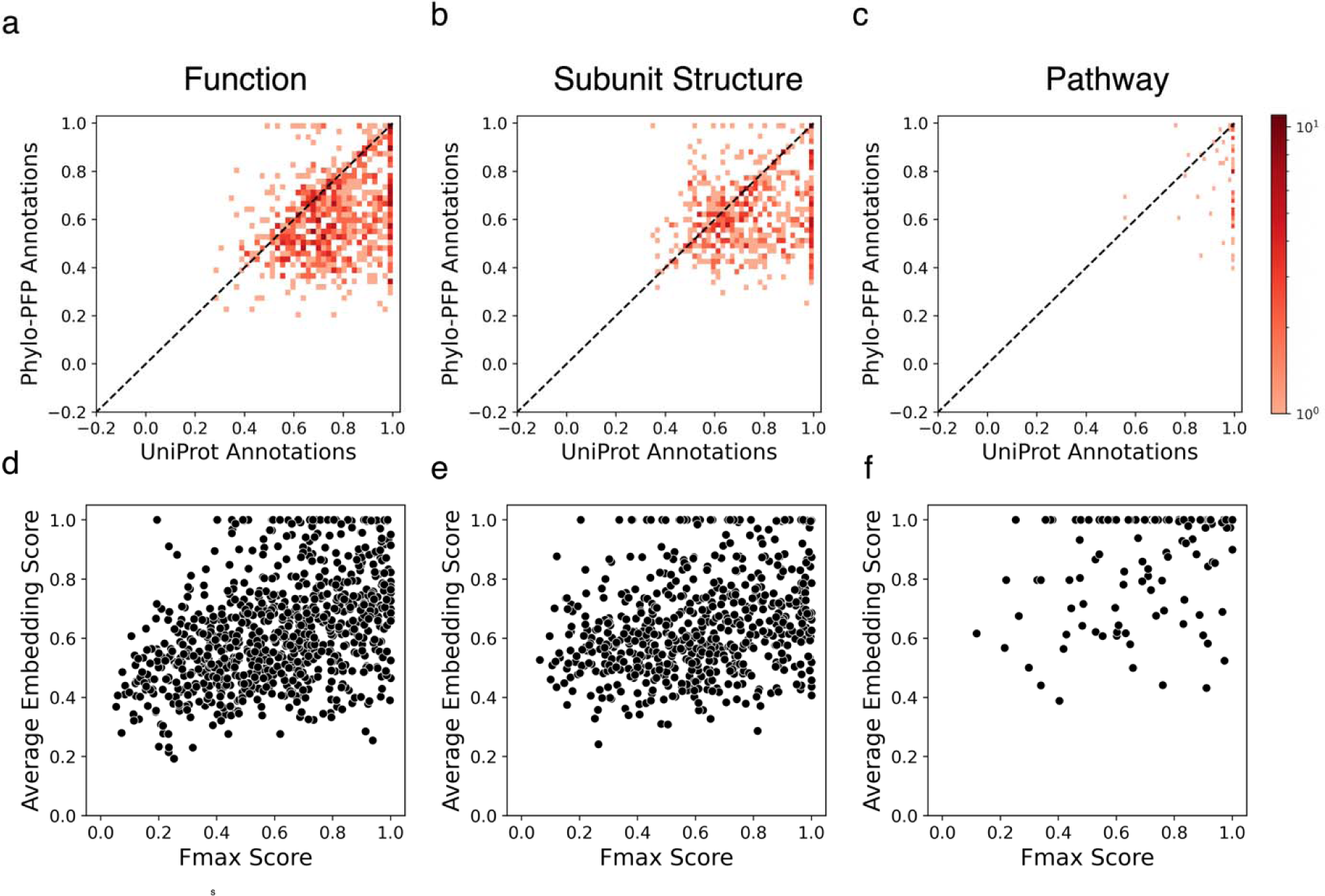
Average embedding scores of predicted GO terms. Predictions were produced by Phylo-PFP. 843, 116, and 632 proteins were used for Function (panel a and d), Subunit structure (b and e), and Pathway (c and f) paragraphs, respectively. a, b, c, the average embedding scores of summaries computed for predicted GO terms relative to the ground truth GO terms taken from UniProt. The color scale shows the number of cases at each point. d, e, f, the average embedding score relative to Fmax score, which indicates the accuracy of predicted GO terms by Phylo-PFP.

Fig. 5d, e, f depict the correlation between the average embedding score and the function prediction accuracy, as measured by the Fmax score^48^. Fmax ranges from 0 to 1, with 1 for the perfect agreement with the ground truth GO terms. The average embedding score exhibits moderate to weak correlation with the Fmax score of GO terms, with the Pearson correlation coefficient of 0.44 for Function, 0.28 for Subunit, and 0.28 for Pathway. Of greater significance is the observation that even relatively low-scoring GO predictions, with an Fmax score around 0.2, can achieve a moderate average embedding score of 0.5. This indicates that GO2Sum can provide readable and useful summaries even for low-scoring GO predictions. According to CAFA challenges^22,49^ recent prediction methods achieve an Fmax score of about 0.5 to 0.6 on average, which is sufficient for GO2Sum to produce meaningful summary. For example, when the Fmax score is 0.5 or higher, 81.9% of proteins have a summary with an average embedding score of 0.5 or higher in Function paragraphs. Similarly, for Subunit Structure and Pathway, the figures are 82.8% and 96.7%, respectively.

## Discussion

We have developed GO2Sum, a fine-tuned variant of the T5 model designed specifically for summarizing the descriptions of GO terms into Function, Subunit, and Pathway paragraphs in UniProt entries. The summaries generated by GO2Sum demonstrated a significantly higher level of agreement with UniProt data when compared to the results obtained using the standard T5 model. Our evaluation of the agreement with the ground truth UniProt paragraphs primarily relied on embedding-based scoring methods, which exhibited a strong correlation with assessments made by human evaluators.

While GO2Sum generally produces accurate summaries, a notable limitation is its inability to mention specific protein names. This limitation arises from the fact that GO descriptions typically lack such specific details. In contrast, UniProt’s Subunit paragraphs often contain specific protein names. To address this limitation and enable the model to provide specific information, it would require access to additional information sources, such as protein-protein interaction data. Integrating GO terms with these external sources represents an intriguing avenue for future research. Another potential future improvement is enhancing the model’s ability to discern which part of a summary corresponds to each GO term description, possibly by implementing a Question and Answering framework^50,51^.

For many years, protein function prediction has relied on GO terms to describe biological functions. However, recent advancements in language models have allowed us to translate these predictions into human-readable text, providing a more intuitive and user-friendly approach to describing predicted functions. We believe that this development contributes significantly to enhancing the interaction between humans and machine learning models, particularly in the context of assisting biologists and medical scientists in their daily research endeavors.

## Acknowledgment

This work was partly supported by the National Science Foundation (DBI2003635, DBI2146026, IIS2211598, DMS2151678, CMMI1825941, and MCB1925643) and by the National Institutes of Health (R01GM133840, 3R01GM133840-02S1).

## Availability

The source code of GO2Sum is available at https://github.com/kiharalab/GO2Sum.

