## Supplementary for "GO2Sum: Generating Human Readable Functional Summary of Proteins from GO Terms"

^1^ Department of Computer Science, Purdue University, West Lafayette, Indiana, 47907, USA

^2^ Department of Biological Sciences, Purdue University, West Lafayette, Indiana, 47907, USA

**Supplementary Information 1.**

We provide three examples of functional descriptions in SwissProt.

Protein:	A9AJN2 **(Phosphatidylserine decarboxylase proenzyme)**

Function:	Catalyzes the formation of phosphatidylethanolamine from phosphatidylserine.

Subunit:	Heterodimer of a large membrane-associated beta subunit and a small pyruvoyl-containing.

Pathway:	Phospholipid metabolism; phosphatidylethanolamine biosynthesis; phosphatidylethanolamine from CDP-diacylglycerol: step 2/2.

Protein:	Q99PM9 (Uridine-cytidine kinase 2)

Function:	Phosphorylates uridine and cytidine to uridine monophosphate and cytidine monophosphate. Does not phosphorylate deoxyribonucleosides or purine ribonucleosides. Can use ATP or GTP as a phosphate donor.

Subunit:	Homotetramer.

Pathway:	Pyrimidine metabolism; CTP biosynthesis via salvage pathway; CTP from cytidine: step 1/3. ; PATHWAY: Pyrimidine metabolism; UMP biosynthesis via salvage pathway; UMP from uridine: step 1/1.

Protein:	O00762 (Ubiquitin-conjugating enzyme E2 C)

Function:	Accepts ubiquitin from the E1 complex and catalyzes its covalent attachment to other proteins. In vitro catalyzes 'Lys-11'- and 'Lys-48'-linked polyubiquitination. Acts as an essential factor of the anaphase promoting complex/cyclosome , a cell cycle-regulated ubiquitin ligase that controls progression through mitosis. Acts by initiating 'Lys-11'-linked polyubiquitin chains on APC/C substrates, leading to the degradation of APC/C substrates by the proteasome and promoting mitotic exit.

Subunit:	Component of the APC/C complex, composed of at least 14 distinct subunits that assemble into a complex of at least 19 chains with a combined molecular mass of around 1.2 MDa. Within this complex, directly interacts with ANAPC2.

Pathway:	Protein modification; protein ubiquitination.

**Supplementary Figure 1. Training and Validation Loss**

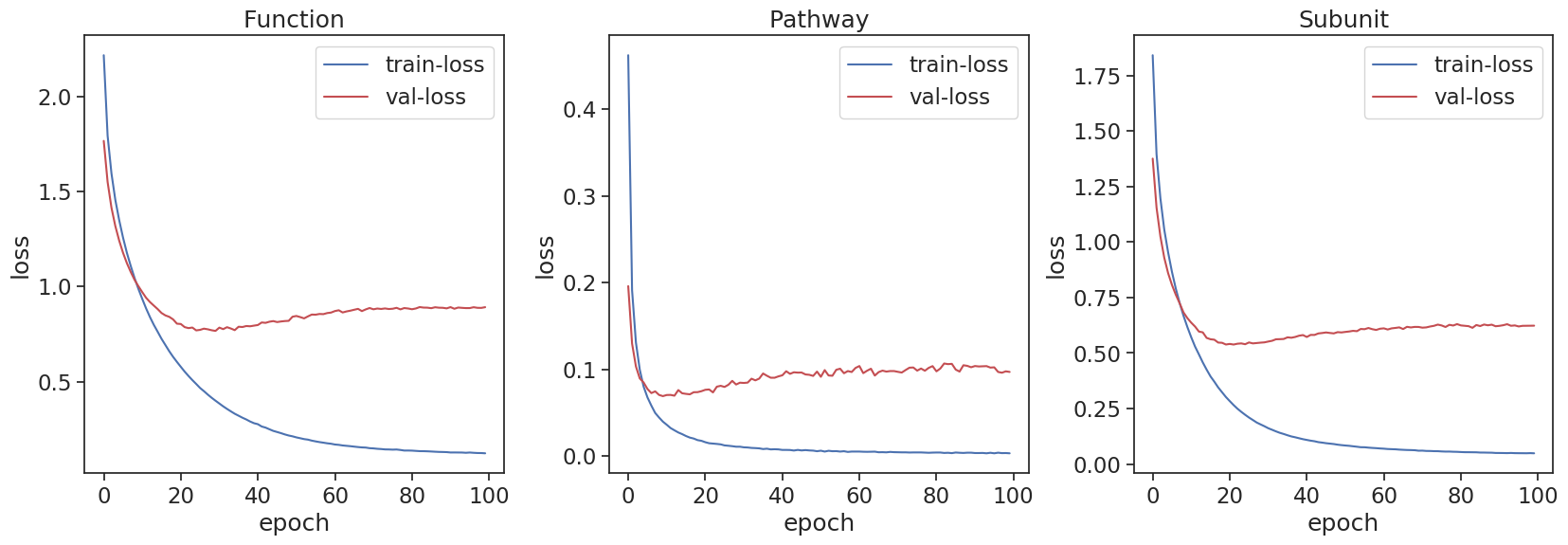

We continued training up to 100 epochs to prevent overfitting and explore parameters even after achieving the best model with the least validation and training loss. Cross-entropy loss function was used in the finetuning process. The fine-tuning process was stopped after the model loss no longer decreased, and the models were trained using NVIDIA RTX5500 GPU with 24GB memory. The best models were achieved for Pathway in 22 epochs, Subunit in 36 epochs, and Function in 34 epochs.

**Supplementary Figure 2. Pearson Correlation Coefficients between BERT and MINILM, and BERT and BioSentVec for Function**

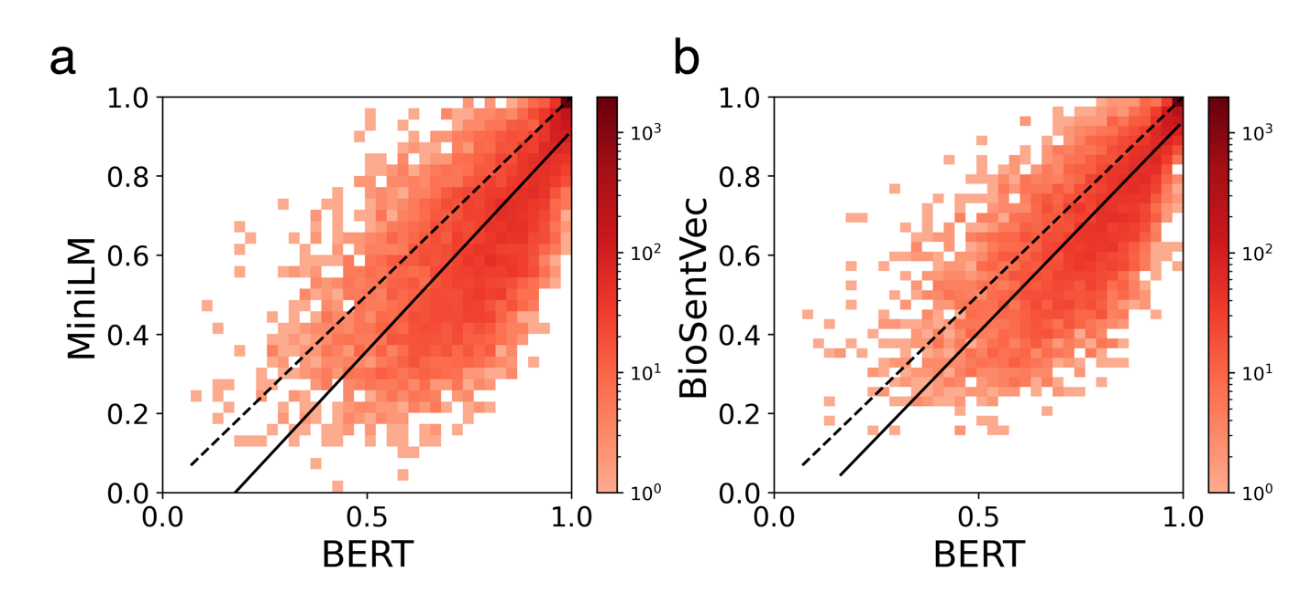

We computed the Pearson correlation coefficient for BERT and MINILM as well as BERT and BioSentVec. The resulting correlation coefficient between BERT and MINILM was 0.79, while the correlation coefficient between BERT and BioSentVec was 0.83.

**Supplementary Figure 3. Embedding scores for BERT, MiniLM, and BioSentVec for Subunit Substructure**

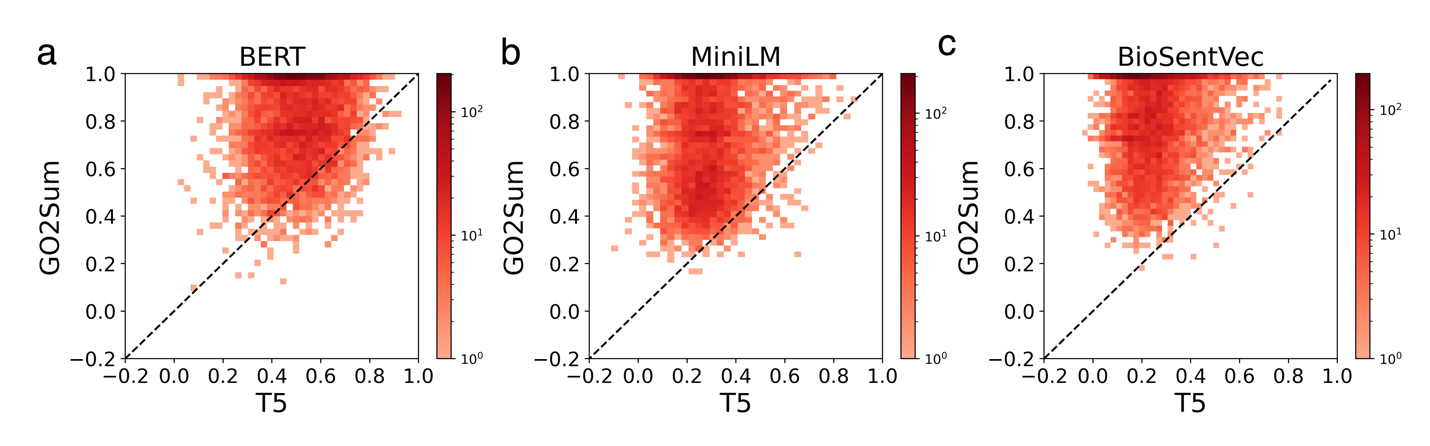

The performance of the model on the Subunit test set using three different embedding-based scores: BERT, MiniLM, and BioSentVec. The results show that GO2Sum performs better than T5(without fine-tuning) for all three scores, with the highest improvement observed for BioSentVec, where the model outperforms T5 in 99.6% of the entries. For MiniLM and BERT, the model performs better than T5 at 98.2% and 92.9% of the entries, respectively.

**Supplementary Figure 4. Embedding scores for BERT, MiniLM, and BioSentVec for Pathway**

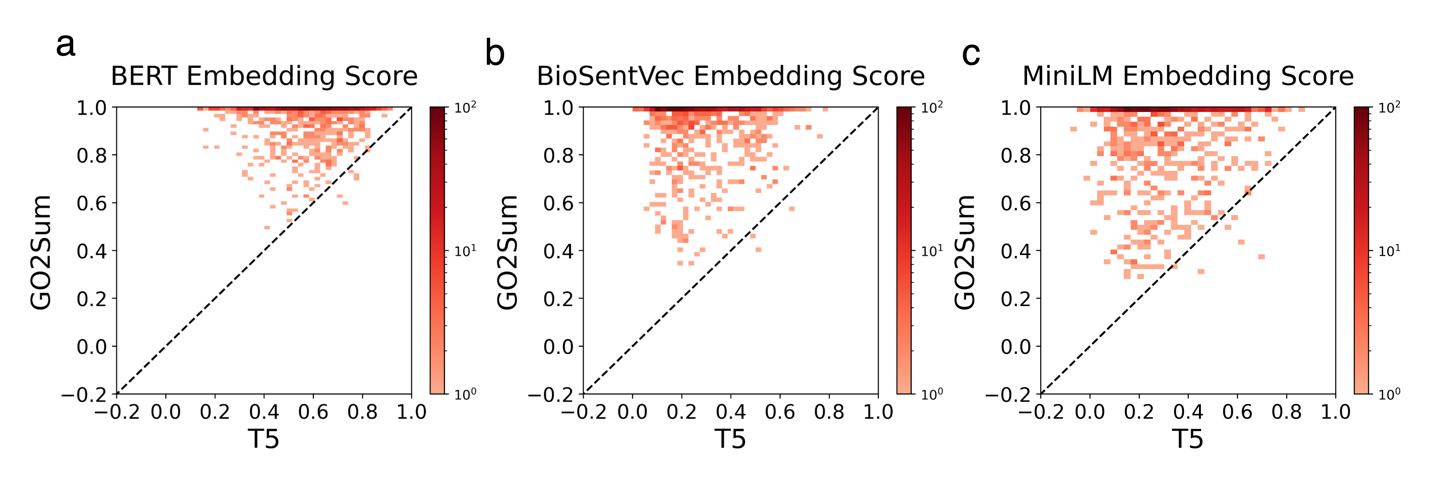

The performance of the model on the Subunit test set using three different embedding-based scores: BERT, MiniLM, and BioSentVec. The results show that GO2Sum performs better than T5(without fine-tuning) for all three scores, with the highest improvement observed for MiniLM, where the model outperforms T5 in 99.6% of the entries. For BioSentVec and BERT, the model performs better than T5 at 99.2% and 92.2% of the entries, respectively.

**Supplementary Figure 5. Pearson Correlation Coefficients between BERT and MINILM, and BERT and BioSentVec for Pathway**

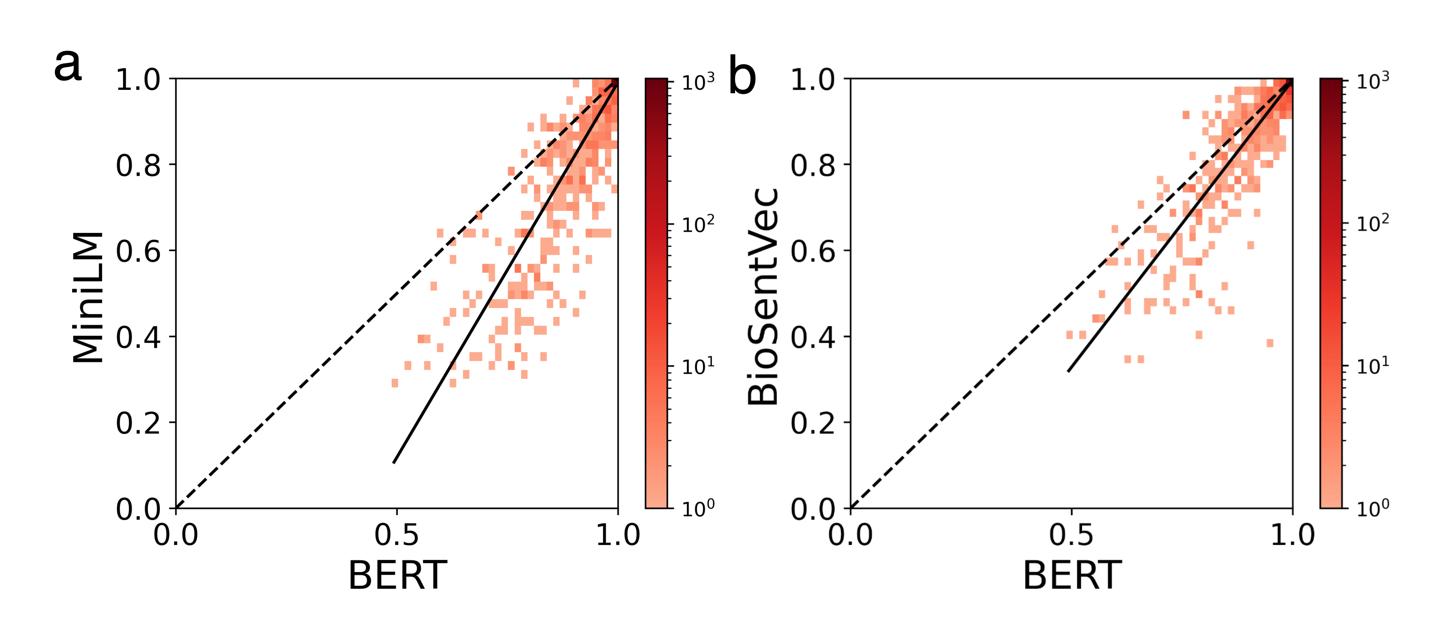

The correlation between the embedding scores, we computed the Pearson correlation coefficient for BERT and MINILM as well as BERT and BioSentVec. The resulting correlation coefficient between BERT and MINILM was 0.90 while the correlation coefficient between BERT and BioSentVec was 0.91.

**Supplementary Information 2. Manual for Human Evaluation Section**

The document below was provided to the six human evaluators for instruction.

**---**

Read the GO Description and UniProt Description, and rate the summaries generated by GO2Sum.

Score 1- 4 (1- Lowest, 4- Highest) and also explain the reason of the score in a line or two.

Criteria of Score 4: Covers all major protein functions and protein name correctly, conveys similar paragraph as UniProt (wording may be different but meaning is the same). Should not contain any wrong protein function or wrong protein name.

Example:

Q9P7X6

GO Description:

A heterotetrameric protein complex that associates with replication origins, where it is required for the initiation of DNA replication, and with replication forks. A membrane-bounded organelle of eukaryotic cells in which chromosomes are housed and replicated. In most cells, the nucleus contains all of the cell's chromosomes except the organellar chromosomes, and is the site of RNA synthesis and processing. In some species, or in specialized cell types, RNA metabolism or DNA replication may be absent. The part of the cytoplasm that does not contain organelles but which does contain other particulate matter, such as protein complexes. The Y-shaped region of a nuclear replicating DNA molecule, resulting from the separation of the DNA strands and in which the synthesis of new strands takes place. Also includes associated protein complexes. Any DNA replication initiation involved in mitotic cell cycle DNA replication. Any DNA strand elongation involved in mitotic cell cycle DNA replication.

UniProt: FUNCTION:

The GINS complex plays an essential role in the initiation of DNA replication.

GO2Sum’s Summary:

The GINS complex plays an essential role in the initiation of DNA replication.

Reason of the Score:

Covers all the Protein function and Protein names correctly.

Criteria of Score 3: Covers most (70% and above) of the protein's functions, but there may be minor differences or omissions regarding some specific protein functions, protein name compared to UniProt's paragraph. Should not contain any wrong protein function or wrong protein name.

Example:

Q54CZ2

GO Description:

A semiautonomous, self replicating organelle that occurs in varying numbers, shapes, and sizes in the cytoplasm of virtually all eukaryotic cells. It is notably the site of tissue respiration. The inner, i.e. lumen-facing, lipid bilayer of the mitochondrial envelope. It is highly folded to form cristae. The chemical reactions and pathways resulting in the formation of ubiquinone, a lipid-soluble electron-transporting coenzyme. The component of mitochondrial inner membrane consisting of gene products and protein complexes that are loosely bound to one of its surfaces, but not integrated into the hydrophobic region.

UniProt FUNCTION:

Component of the coenzyme Q biosynthetic pathway. May play a role in organizing a multi-subunit COQ enzyme complex required for coenzyme Q biosynthesis. Required for steady-state levels of other COQ polypeptides.

GO2Sum’s Summary:

Component of the coenzyme Q biosynthetic pathway. May play a role in organizing a multi-subunit COQ enzyme complex required for coenzyme Q biosynthesis

Reason of the Score:

The two descriptions share the common function of being a component of the coenzyme Q biosynthetic pathway and potentially organizing a multi-subunit COQ enzyme complex. However, the UniProt description adds the additional aspect of the protein's requirement for steady-state levels of other COQ polypeptides.

Criteria of Score 2: Partially covers the protein's function but lacks some important details and there may be major differences or omissions regarding some specific protein functions, protein name compared to UniProt's paragraph. May contain some wrong protein name or wrong protein function.

Example:

Q05783

GO Description:

Any molecular function by which a gene product interacts selectively and non-covalently with DNA	Binding to a DNA structure formed by the minor groove of adenine-thymine-rich DNA regions. Examples of proteins having this function are AT-rich interaction domain -containing proteins. A membrane-bounded organelle of eukaryotic cells in which chromosomes are housed and replicated. In most cells, the nucleus contains all of the cell's chromosomes except the organellar chromosomes, and is the site of RNA synthesis and processing. In some species, or in specialized cell types, RNA metabolism or DNA replication may be absent. A structure composed of a very long molecule of DNA and associated proteins	that carries hereditary information. Any process that results in the specification, formation or maintenance of the physical structure of eukaryotic chromatin. Any process that modulates the frequency, rate or extent of transcription mediated by RNA polymerase II. The activity of binding selectively and non-covalently to and distorting the original structure of DNA, typically a straight helix, into a bend, or increasing the bend if the original structure was intrinsically bent due to its sequence.

UniProt FUNCTION:

Binds preferentially single-stranded DNA and unwinds double-stranded DNA. Prefers sites containing the sequence 5'-ttg-3'. Facilitates DNA bending. Associated with early embryonic chromatin in the absence of histone H1.

GO2Sum’s Summary:

Binds preferentially AT-rich DNA sequences.

Reason of the Score:

The GO2Sum description lacks important details mentioned in the UniProt paragraph, such as the protein's ability to bind and unwind DNA, its preference for specific DNA sequences, and its association with early embryonic chromatin. These differences and omissions regarding important protein functions justify a score of 1, indicating that the GO2Sum description partially covers the protein's function but lacks significant details.

Criteria of Score 1: Totally Irrelevant. UniProt does not match GO2Sum summary and is totally irrelevant.

Example:

P27473

GO Description:

The contents of a cell excluding the plasma membrane and nucleus, but including other subcellular structures. Any immune system process that functions in the calibrated response of an organism to a potential internal or invasive threat. Any process that results in a change in state or activity of a cell or an organism as a result of a stimulus from a bacterium.

UniProt FUNCTION:

This protein aggregates to form microtubular structures.

GO2Sum’s Summary:

Required for the anti Gram-positive activity of B.anthracis.

Reason of the Score:

Based on the provided function descriptions, the functions mentioned in UniProt and GOSum appear to be unrelated or irrelevant to each other. The UniProt function describes the protein's involvement in forming microtubular structures, while the GOSum function relates to its role in the anti Gram-positive activity of B.anthracis. These functions are distinct and do not seem to overlap or have a direct connection.

**Supplementary Table 1. Examples of Summary generated for UniProt Subunit.**

| **Protein Name** | **ID** | **UniProt** | **T5** | **GO2Sum** | **Average embedding score** |
| --- | --- | --- | --- | --- | --- |
| H/ACA ribonucleoprotein complex subunit 2-like protein | Q9LEY9 | Component of the small nucleolar ribonucleoprotein particle containing H/ACA-type snoRNAs | molecule. Binding to messenger RNA (mRNA). Binding to | Component of the small nucleolar ribonucleoprotein particles containing H/ACA-type snoRNAs | 0.98 |
| Apolipoprotein M | O95445 | Interacts with LRP2; LRP2 mediates APOM renal uptake and subsequent lysosomal degradation. | a phospholipid bilayer. a phospholipid bilayer. | Interacts with LRP2; LRP2 mediates APOM renal uptake and subsequent degradation. | 0.96 |
| Guanine nucleotide-binding protein G(o) subunit alpha | P59215 | Interacts with RGS14. G proteins are composed of 3 units; alpha, beta and gamma. The alpha chain contains the guanine nucleotide binding site. | the receptor. Binding to a cyclic AMP receptor. Bind | G proteins are composed of 3 units; alpha, beta and gamma. The alpha chain contains the guanine nucleotide binding site. | 0.93 |
| Actin-related protein 5 | P53946 | Component of the chromatin-remodeling INO80 complex, at least composed of ARP4, ARP5, ARP8, RVB1, RVB2, TAF14, NHP10, IES1, IES3, IES4, IES6, ACT1, IES2, IES5 and INO80. | the nucleus. the nucleus. chromosomes. | Component of the chromatin-remodeling INO80 complex. | 0.86 |
| Nucleoside diphosphate kinase 1 | B4FK49 | Homohexamer. Can also form dodecamers. | ATP. cytidine (5'-) triphosphate. | Homohexamer. | 0.79 |
| Serine/threonine-protein kinase 4 | A4K2W5 | Homodimer; mediated via the coiled-coil region. Interacts with NORE1, which inhibits autoactivation. Interacts with and stabilizes SAV1. Interacts with RASSF1. Interacts with FOXO3. Interacts with RASSF2 Interacts with AR, PKB/AKT1, TNNI3 and SIRT1. Interacts with DLG5 Interacts with MARK3 and SCRIB in the presence of DLG5. | the organellar chromosomes. the nucleus is the site. | Homodimer; mediated via the coiled-coil region. | 0.58 |
| **NAD(P)H dehydrogenase [quinone] 1** | P15559 | Homodimer. Interacts with PDLIM4 isoform 2; this interaction stabilizes PDLIM4 isoform 2 in response to oxidative stress and protects it from ubiquitin-independent degradation by the core 20S proteasome. Interacts with TP73 ; this interaction is NADH-dependent, stabilizes TP73 in response to oxidative stress and protects it from ubiquitin-independent degradation by the 20S proteasome Interacts with TP53; this interaction is NADH-dependent, stabilizes TP53 in response to oxidative stress and protects it from ubiquitin-independent degradation by the 20S proteasome | the cytoplasm, but not the cytoplasm. Any process that results in | Homodimer. | 0.36 |
| Protein OBERON 2 | Q9LUB7 | Self-interacts Interacts with OBE1, OBE3 and OBE4 Binds to VPg of pea seed borne mosaic virus , turnip mosaic virus and lettuce mosaic virus , but not with VPg of tobacco etch virus , cowpea mosaic virus , tomato black ring virus and grapevine fan leaf virus. | the nucleus contains all of the cell's chromosomes except the. | Homodimer. | 0.22 |

Q9LEY9, O95445, and P59215 had high average embedding based-score ranging from 0.93 - 0.98. The scores of Q9LEY9 and O95445 were 0.98 and 0.96, respectively. The information in both summaries was almost identical to the ground truth, with only minor differences. For P59215, GO2Sum had an average embedding score of 0.93. Here the model's output was not as complete as the ground truth. The main statement in both the ground truth and the GO2Sum's output was the same, but the model's output did not include additional information about “Interacts with RGS14”. T5 generated summary were wrong and none of them generated the Subunit paragraphs correctly.

The next three examples, P53946, B4FK49, and A4K2W5, are cases where GO2Sum had a moderate average embedding-based score between 0.58 and 0.86. These two proteins have incomplete Subunit structure descriptions. The summaries by GO2Sum correctly captured some Subunit structure of the proteins but lacked some of them. T5 again gave fragmented phrases.

The last two proteins, P15559 and Q9LUB7, received low average embedding scores. P15559 had an average embedding score of 0.36, and Q9LUB7 had an average embedding score of 0.22. By looking at the Subunit structure description paragraph of these proteins, we can see that the descriptions are very detailed and contain a lot of specific protein names. In contrast, the GO descriptions are more generic. This is the reason why these proteins received low scores, as it had difficulty capturing the specific protein names from the generic GO descriptions.

**Supplementary Table 2. Examples of Summary generated for UniProt Pathway.**

| **Protein Name** | **ID** | **UniProt** | **T5** | **GO2Sum** | **Average embedding score** |
| --- | --- | --- | --- | --- | --- |
| Spermidine N(1)-acetyltransferase | P0DKR8 | Amine and polyamine degradation; spermidine degradation. Amine and polyamine degradation; spermine degradation. | a magnesium (Mg) ion. Catalysis of the reaction. | Amine and polyamine degradation; spermine degradation. | 0.98 |
| Succinate dehydrogenase [ubiquinone] iron-sulfur subunit, mitochondrial | Q09545 | Carbohydrate metabolism; tricarboxylic acid cycle; fumarate from succinate: step 1/1. | the ubiquinone cycle with the respiratory chain. the ubiquinon | Carbohydrate metabolism; tricarboxylic acid cycle; fumarate from succinate. | 0.95 |
| Mitochondrial ubiquitin ligase activator of NFKB 1 | Q8VCM5 | Protein modification; protein ubiquitination. ; PATHWAY: Protein modification; protein sumoylation. | ubiquitin. Y-SUMO + X, where both | Protein modification; protein ubiquitination. | 0.92 |
| Cholesterol side-chain cleavage enzyme, mitochondrial | Q9QZ82 | Lipid metabolism; C21-steroid hormone metabolism. Steroid metabolism; cholesterol metabolism. | heme. The chemical reactions and pathways resulting in the formation of steroid | Lipid metabolism; steroid biosynthesis. | 0.84 |
| Methionine synthase | O33465 | Amino-acid biosynthesis; L-methionine biosynthesis via de novo pathway; L-methionine from L-homocysteine: step 1/1. | cobalt atom. a cobalt atom. a | Amino-acid biosynthesis. | 0.73 |
| NADH:FAD oxidoreductase | O87008 | Xenobiotic degradation. | protein. a protein. a molecule. a | Aromatic compound metabolism. | 0.55 |
| Putative mRNA-capping enzyme P5 | P14583 | mRNA processing; mRNA capping. | guanosine triphosphate. Catalysis of the reaction. | Purine metabolism; 7-cyano-7-deazaguanine biosynthesis. | 0.39 |

The first three examples (UniProt ID: P0DKR8, Q09545, and Q8VCM5) are cases where GO2Sum had the average embedding-based score of 0.92 to 0.98. With these high scores, GO2Sum made an almost perfect summary to the ground truth with a subtle difference.

For P0DKR8, the same pathway details are repeated in the ground truth in UniProt, where UniProt associated a different evidence code to each of them. GO2Sum was correct in eliminating the repetitive information. In Q09545, GO2Sum failed to capture the step information and in Q8VCM5 it failed to capture one of the pathways. T5 again gave fragmented phrases. Furthermore, it is worth noting that the vanilla T5 model's generated sentences, which did not include any correct pathway information.

Moving on to the next two examples, Q9QZ82 and O33465, where GO2Sum had a moderate average embedding-based score ranging from 0.73 to 0.84. In these cases, GO2Sum managed to capture some pathway information, but it also missed some details. For example, in the case of Q9QZ82, GO2Sum failed to capture the pathway information related to "Steroid metabolism" and "Cholesterol metabolism." Similarly, for O33465, GO2Sum missed out on summarizing the pathways "L-methionine biosynthesis via de novo pathway" and "L-methionine from L-homocysteine," as well as the step information associated with these pathways.

The summary provided by GO2Sum received low scores of 0.55 and 0.39 for O87008 and P14583, respectively. In fact, a score of 0.39 was the lowest among the pathway paragraphs GO2Sum generated. Interestingly, GO2Sum generated different pathways compared to UniProt. For O87008, UniProt wrote "Xenobiotic degradation", while GO2Sum indicated "Aromatic compound metabolism." Similarly, UniProt categorized P14583 as involved in "mRNA processing; mRNA capping," but GO2Sum suggested "Purine metabolism; 7-cyano-7-deazaguanine biosynthesis." When we took a closer look at the GO Description for these proteins, we realized that the descriptions were too general without a mention of more specific pathways.
